# Serial dependence in haptic perception during active manual exploration

**DOI:** 10.64898/2026.09.23.753977

**Authors:** Yunshan Zhang, Tatsuhiro Nakamura, Takashi Hanakawa, Tatsuya Umeda

## Abstract

Serial dependence, whereby current perception is systematically biased toward the most recent sensory experience, has been demonstrated across multiple sensory modalities. However, whether serial dependence occurs during active haptic perception remains unknown. Moreover, because haptic perception of object geometry depends on hand movements for sensory acquisition (i.e., active touch), it remains unclear whether serial dependence, if any, originates from perceptual processing, movement, or both. Here, we investigated whether active haptic perception exhibits attractive serial dependence using a newly developed haptic device that let participants explore virtual slopes of different heights. Participants reported perceived slope heights, and we recorded their kinematics. Perceptual judgments were systematically attracted toward the most recently experienced slope, demonstrating serial dependence during active haptic perception. By contrast, none of the kinematic parameters exhibited corresponding serial dependence. These findings show that serial dependence extends to the somatosensory perceptual domain and suggest that the observed perceptual bias was not mediated by systematic changes in hand movements.

## Introduction

Humans perceive the geometry of objects, such as their size and shape, by actively exploring their surfaces with the hand. This ability, known as haptic perception, relies on integrating tactile signals conveying local surface features with proprioceptive signals conveying hand position and movement (Berryman et al., 2006; Hsiao, 2008; Sobinov and Bensmaia, 2021; Voisin et al., 2002). By combining these complementary sources of somatosensory information, humans can estimate object geometry with remarkable precision, often approaching the performance achieved through vision (Henriques and Soechting, 2003).

Perception is not determined solely by incoming sensory input. Instead, the brain continuously integrates current sensory evidence with prior information derived from long-term environmental statistics (Stevens and Greenbaum, 1966; Weiss et al., 2002), explicit expectations (Kok et al., 2012), and recent sensory history (Fischer and Whitney, 2014; Vogels et al., 1996). This concept is commonly described within a Bayesian framework, in which prior knowledge and noisy sensory input are combined to generate stable perceptual estimates (Kersten et al., 2004; Petzschner et al., 2015; Summerfield and de Lange, 2014). Among these sources of prior information, recent sensory experience exerts a particularly strong influence on perception. One well-known example in haptic perception is the curvature aftereffect, in which prolonged exploration of a curved surface induces a subsequent repulsive perceptual bias (Vogels et al., 1996). Another prominent history-dependent phenomenon is serial dependence, whereby perception of the current stimulus is systematically attracted toward recently experienced stimuli (Cicchini et al., 2024; Manassi et al., 2023; Pascucci et al., 2023). Serial dependence has been demonstrated across a broad range of visual tasks, from orientation estimation (Fischer and Whitney, 2014; Fritsche et al., 2017), to high-level judgments such as face recognition (Liberman et al., 2014; Manassi and Whitney, 2022), numerosity estimation (Cicchini et al., 2014), and attractiveness evaluation (Chang et al., 2017; Kim et al., 2019), and has also been reported in audition and olfaction (Burg et al., 2022; Motala et al., 2020). Although researchers have extensively investigated serial dependence in multiple sensory modalities, it remains unknown whether haptic perception in the motor-somatosensory domain also exhibits attractive serial dependence.

Haptic perception is unique in that sensory information about object geometry is acquired through self-generated exploratory movements (Delhaye et al., 2018). Because perceptual judgments are inherently coupled with behavior, movement itself may change over time under the influence of sensory stimuli. This raises the possibility that any serial dependence observed during active touch could reflect changes in perceptual processing, behavior, or both. However, previous studies of serial dependence have not directly distinguished between these possibilities.

In the present study, we addressed this question using a newly developed haptic device that enabled participants to actively explore virtual slopes of precisely controlled heights in a two-dimensional workspace. Participants reported perceived slope height after each trial, while we recorded multiple kinematic parameters simultaneously. We show that perceived slope height exhibits robust serial dependence, whereas none of the quantified kinematic parameters during movements display corresponding serial dependence. These findings extend serial dependence to active haptic perception and indicate that the observed bias is more likely to arise from perceptual processing than from the quantified kinematics.

## Methods

### Participants

Twenty-three right-handed volunteers [mean age = 22.7 years, standard deviation (SD) = 3.4, age range = 19–31; 9 females, 14 males] participated in Experiment 1, in which they estimated the height of virtual slopes. Among them, 21 participants (mean age = 22.9 years, SD = 3.5, age range = 19–31; 8 females, 13 males) also completed Experiment 2 to measure discrimination thresholds. The sample size was comparable to those used in previous studies of visual serial dependence^14^. Participants were naïve to the purpose of the experiment and reported no history of somatosensory, neurological, or psychiatric disorders. We obtained written informed consent from all participants before the experiments. The Kyoto University Graduate School and Faculty of Medicine Ethics Committee approved the study (C1624-1), and it was conducted in accordance with the Declaration of Helsinki.

### Apparatus

A custom-built manipulandum consisted of a horizontally oriented handle driven by the Touch USB (3D Systems Inc.) (Fig. 1a). The handle (6 mm in diameter and 72 mm in length) was attached to the stylus and movable only in a two-dimensional (2D) plane perpendicular to the ground using two linear ball slides (LS1052, LSP25150, THK Co. Ltd.). Participants grasped the proximal end of the handle with the right thumb and index finger (Fig. 1a). The range of movements was 93 mm on the horizontal axis and 24.8 mm on the vertical axis. A counterweight compensated for part of the handle weight to prevent overheating during repeated movements.

**Figure 1.**
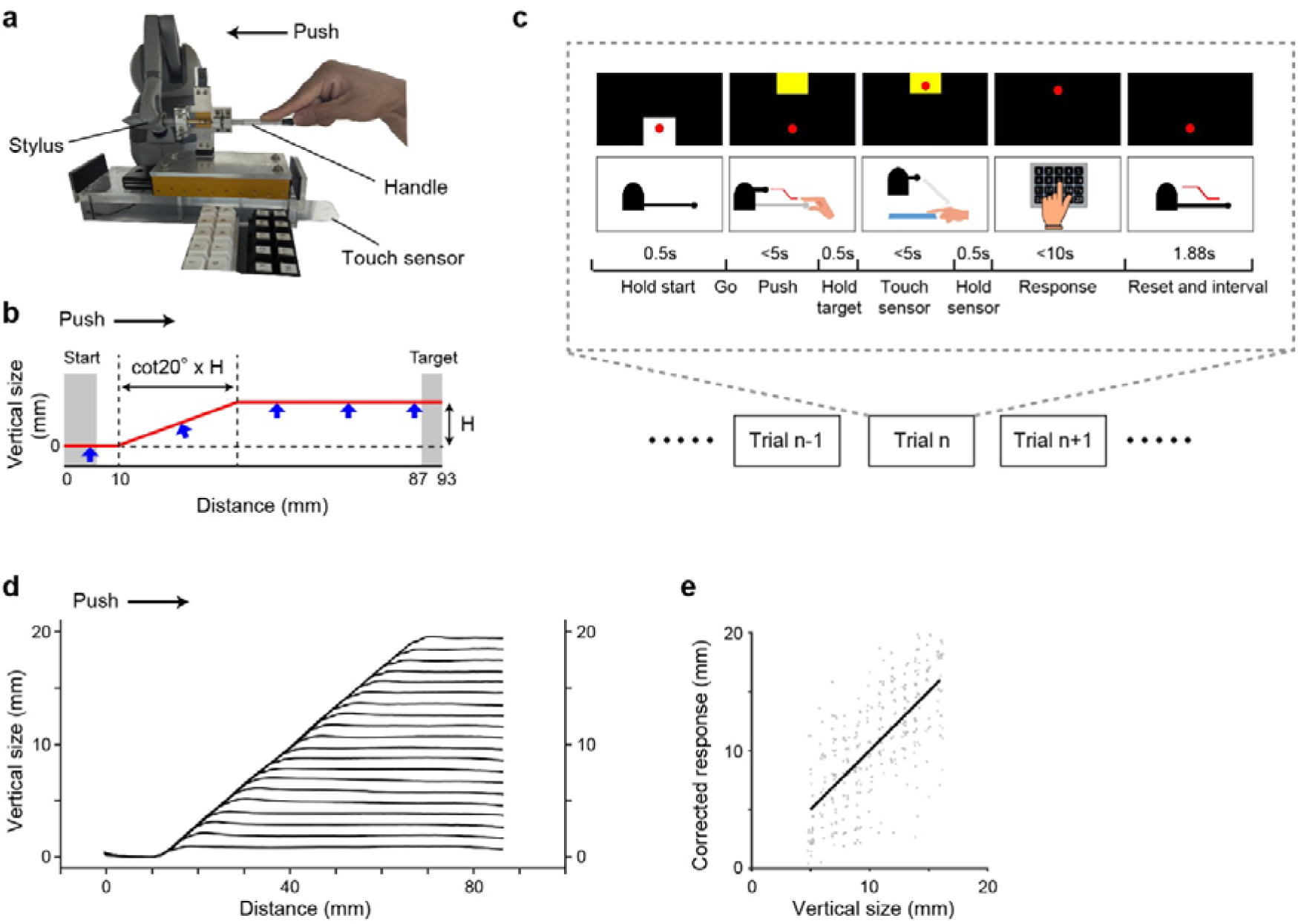
Experimental setup and behavioral performance. **a** Experimental apparatus. **b** Schematic illustration of the virtual slope presented by the haptic device. A 20-degree slope starts 10 mm from the edge of a linear ball slide and rises to a height of 1-20 mm (denoted as H). The red line indicates the boundary of the virtual slope, and blue arrows indicate the resistive force generated when the distal end of the stylus crossed the slope boundary. **c** Experimental paradigm. Participants pushed the handle to explore randomly presented virtual slopes (1–20 mm in 1-mm increments) and reported the perceived slope height in each trial by pressing one of 20 response buttons. On the monitor, the red cursor indicated the handle position, whereas the white and yellow squares indicated the start and target positions, respectively. **d** Mean vertical handle trajectories for the 20 slope heights from a representative participant. Trajectories were aligned to the proximal edge of the linear ball slide. Shaded areas indicate SEM. **e** Relationship between presented and reported slope heights after correction for central tendency and overall response bias in a representative participant. The solid line indicates the linear regression fit. Overlapping data points are displayed with slight offsets for visualization.

The haptic system presented a virtual slope with one of 20 heights (1–20 mm in 1-mm increments) (Fig. 1b). After the initial horizontal straight path (from x=0 mm, y=0 mm to x=10 mm, y=0 mm), the slope emerged with an inclination of 20 degrees (shown as a red line in Fig. 1b). When the distal end of the stylus crossed the boundary of the slope, the resistance was generated by a force perpendicular to the slope boundary and proportional to the penetration depth with a stiffness of 1 N/mm. Resultantly, when the participants moved the manipulandum, they perceived it as if they were stroking the surface of an object with the stylus. When the slope reached a predetermined height y=H mm, the path turned to a horizontal line to x=87 mm, y=H mm. The manipulandum was controlled by custom-written programs for Touch USB control, and the task procedure was controlled by MonkeyLogic (National Institute of Mental Health) (Hwang et al., 2019). The two programs were synchronized with each other via digital signals. They controlled the direction, timing, and strength of the force applied to the stylus. Device encoders measured the position and velocity of the manipulandum movement.

The participants sat facing the workspace, ensuring that the virtual slopes were presented within the plane that included both their body’s midline and the manipulandum’s movable space. The manipulandum was positioned under the table, making the handle invisible to the participants. A liquid crystal display monitor displaying the back-forward position of the handle was placed on the table in front of each participant. A touch sensor plate was located between the manipulandum and the participants to keep their right hand at the starting position of each trial. A keyboard was placed next to the manipulandum so that participants could use their left hand to report the perceived height of the virtual slope.

### Task Procedure

In the experiment, the participants were requested to estimate the height of the virtual slope. Each trial began when a white square indicating the start area appeared at the bottom of the monitor (Fig. 1c). A red cursor on the monitor represented the back-forward position of the handle. If the red cursor corresponding to the handle position entered the start area and remained there for 500 ms, the white square would disappear. Instead, a yellow square, signifying the target area, appeared at the top of the monitor, accompanied by a sound cue (2,000 Hz). Upon hearing the start cue, participants pushed the handle to the target area (x = 87 to 93 mm). Participants were required to push the handle to the target within 5 s; otherwise, the trial was considered unsuccessful. When the handle reached the target area and remained there for 500 ms, a double-sound signal (1,000 Hz, 100-ms interval) notified participants that the handle had successfully entered the target area. Participants then released the handle and placed their right hand on the touch sensor plate within 5 s. After touching the sensor for 500 ms, another sound cue (6,000 Hz) prompted participants to report the virtual slope height. They used their left hand to press a key on a keyboard marked from 1 to 20 within 10 s. This report served as a measure of the height participants perceived. After the key was pressed, the system presented another sound signal (3,000 Hz). After a set inter-trial interval (1.88 s), the handle automatically returned to the start area, and the next trial began. Meanwhile, we recorded handle kinematic parameters to quantify hand movements during manual exploration. The experiment consisted of three runs, with 200 trials per run. Trials in which participants failed to complete the task within the time limit were excluded from all analyses.

Before the experimental sessions, participants were familiarized with responding to 20 different virtual slope heights. In the first 10 trials of the training run, only the highest slope (20 mm) was presented, and participants learned how to move the handle along the path and follow the task procedure. Next, they experienced slopes ranging from 1 mm to 20 mm in order and reported their perception. This fixed training run was repeated three times. Finally, as in the main experiment session, participants were randomly presented with virtual slopes with 20 different heights and were asked to report the height they perceived. During the training run, the actual height was displayed on the monitor as feedback after each trial. A total of 200 trials were conducted in this training run.

### Data Analysis

#### Removal of edge artifacts

Responses to a given stimulus are expected to distribute around the presented stimulus value. However, when the stimulus value is located near the upper or lower boundaries of the stimulus range, participants cannot express values beyond the boundaries. In our experimental setting, when the presented stimulus is 20 mm, responses larger than 20 mm were impossible. This restriction distorted the response distribution and artificially biased responses, creating edge artifacts and shifting responses toward the center of the stimulus range.

To minimize the boundary effects, we excluded trials with stimuli near the lower and upper limits of the stimulus range from the analysis. For a normally distributed variable, approximately 10% of observations are expected to exceed μ +1.28σ, where μ and σ denote the mean and standard deviation, respectively. In the pooled response distribution across participants, 1.28σ corresponded to 4.54 mm. Therefore, we included only trials with stimulus heights ranging from 5 mm to 16 mm in the analysis. Within this range, more than 90% of responses were expected to remain within the available response range, allowing the influence of edge artifacts to be considered negligible.

#### Correction of central tendency and overall response biases

Magnitude estimation tasks are known to exhibit central tendency, whereby responses are biased toward the center of the stimulus distribution. As a result, the relationship between presented stimuli and reported responses is typically compressed relative to the identity line. To correct this compression, we fitted a linear regression of presented stimuli on participants’ responses and calculated the slope. We then adjusted responses using the slope, along with the mean stimulus and mean response values.

Beyond central tendency, individual participants also showed an overall response bias, consistently perceiving stimuli as either larger or smaller than they were. To remove these general individual response biases, we demeaned response errors by subtracting each participant’s overall mean response error from each response error. As a result, the regression line had a slope of 1 and passed through the point defined by the mean stimulus and mean response.

The overall correction function we used is

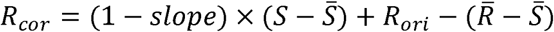

where *R_cor_* represents the corrected response, *R_ori_* represents the original reported response, *S* represents the presented stimulus, 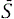 represents the mean of the presented stimulus, and 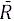 represents the mean of the original responses. The term *slope* represents the regression coefficient obtained from the linear relationship between the original responses and the presented stimuli.

#### Haptic perceptual bias

Response error was defined as the difference between the reported and presented virtual slope height on each trial. To quantify the effects of the previous stimulus on current response, we plotted the response error in the current trial against the relative stimulus difference, defined as the difference between the stimulus presented in the previous trial and that presented in the current trial. In this plot, positive response errors indicate overestimation of the current stimulus, whereas negative values indicate underestimation. Thus, data points with the same sign on the abscissa and ordinate indicate an attractive effect, where the current response was biased toward the previous stimulus. By contrast, data points with opposite signs on the abscissa and ordinate indicate a repulsive effect. We quantified the magnitude of bias toward the previous stimulus using an error-folding approach (Barbosa and Compte, 2020). Specifically, we assigned positive values to response errors indicating attraction toward the previous stimulus, and negative values to response errors indicating repulsion from the previous stimulus. We then calculated the mean of these sign-adjusted response errors and used it as an index of serial dependence.

Next, we subsampled trials to minimize imbalances in both the distribution of relative stimulus differences between successive trials and the number of trials across virtual slope heights. Sufficient trial pairs were available for each relative height difference when the range was restricted to −9 to 9 mm. For each relative stimulus difference, up to five trial pairs were randomly selected; when fewer than five pairs were available, all pairs were retained. We also balanced the trial count across stimulus magnitudes so the difference between the maximum and minimum number of trials did not exceed four. These trial balancing criteria minimized sampling imbalances while preserving the original distribution of stimulus magnitudes (cosine similarity = 0.98). The mean number of subsampled trials was 86.1 ± 2.7.

As a control analysis, we conducted an N+1 analysis to examine whether the apparent serial dependence arose from non-sequential biases in the dataset. Specifically, we tested whether responses in the current trial were influenced by stimuli presented in the next trial. We selected trials using the same procedure described above.

#### Effects of previous stimuli on kinematics

To examine whether previous stimuli systematically influenced motor behavior coupled with haptic perception, we quantified hand movement using five kinematic parameters potentially associated with slope-height perception (Fig. 1b):

(1) mean handle height while moving along the slope, calculated as the time-averaged handle height while the handle was located between x = 10 mm and x = 10 + cot (20**°)** H mm;
(2) mean handle height around the target position, calculated as the time-averaged handle height while the handle was located between x = 10 + cot (20**°)** H mm and x = 87.5 mm;
(3) total force applied to the handle while moving along the slope, calculated as the cumulative force exerted by the device on the handle while the handle was located between x = 10 mm and x = 10 + cot (20**°**) H mm;
(4) duration of movement along the slope, defined as the time elapsed from when the handle reached x = 10 mm until it reached x = 10 + cot (20**°)** H mm; and
(5) movement speed along the slope, calculated as the total distance traveled by the handle between x = 10 mm and x = 10 + cot (20**°)** H mm, divided by the duration of movement along the slope.

The haptic system recorded the position of the stylus attached to the handle and the force exerted on it, together with the corresponding timestamps, at a sampling rate of 1000 Hz.

To remove central tendency and overall biases in kinematic parameters, we applied a correction procedure analogous to that used for perceptual responses:

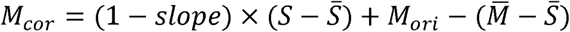

where *M_cor_* represents the corrected kinematic parameters, *M_ori_* represents the original recorded kinematic parameters, *S* represents the presented stimulus, 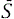 represents the mean stimulus, and 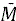 represents the mean of the original kinematic parameters. The term *slope* denotes the regression coefficient obtained from the relationship between the original kinematic parameters and the presented stimulus. To ensure consistency across analyses, we used the same trials selected for the perceptual bias analysis to calculate movement bias.

As with the analysis of sign-adjusted response errors, we examined whether previous stimuli systematically biased kinematics toward previous sensory stimuli. Unlike perceptual responses, however, no predefined expected value exists for a given kinematic parameter at each stimulus height, precluding direct calculation of trial-by-trial errors. Instead, for each kinematic parameter, we first fitted a linear regression model relating the current stimulus to the corrected kinematic values. We then entered the current stimulus for each trial into the fitted model and used the resulting prediction as the expected kinematic value for that trial. We defined the residual difference between the observed and predicted values as the kinematic deviation, representing variability that the current stimulus could not explain.

To further examine whether variations in perceptual judgments were associated with variations in movements within the same trial, we calculated Pearson’s correlation coefficient between perceptual error and the deviation in each of the five kinematic parameters within each participant. This analysis tested whether kinematic variations systematically covaried with perceptual errors within the same participant. We then summarized the resulting correlation coefficients across participants.

### Statistical analysis

We used two-tailed one-sample and paired t-tests, as appropriate. When necessary, the Bonferroni correction was applied to address the family-wise error rate. The significance level (alpha) was set at 0.05. Data are presented as mean ± standard error of the mean (SEM) unless otherwise specified. When data are presented as mean ± standard deviation (SD), this is explicitly indicated. Statistical analyses were performed using MATLAB R2023a (MathWorks). No statistical methods were used to predetermine the sample size; however, sample sizes adhered to published standards.

### Statement of preregistration

This study was not preregistered.

## Results

### Haptic perception of virtual slopes during manual exploration

We developed a haptic system to quantitatively assess the perception of virtual slope height during manual exploration (Fig. 1a–c). Representative handle trajectories from one participant showed distinct height-dependent paths corresponding to the presented virtual slopes (Fig. 1d). Participants correctly identified the presented slope height in 15.3 ± 4.7% (mean ± SD), which was significantly greater than the chance level of 5% (*t*(22) = 10.5, *p* = 5.1 10^-10^, Cohen’s d = 2.2). When responses within ±1 mm of the presented height were considered correct, the accuracy was 39.0 ± 9.7%. Nevertheless, precisely identifying the presented height remained challenging, indicating substantial perceptual uncertainty. Figure 1e shows that perceived slope height scaled with presented slope height after correcting for central tendency and overall response bias, indicating that participants reliably encoded the height differences between virtual slopes.

### Serial dependence in haptic perception

We examined whether haptic perception was systematically influenced by the previous stimulus. To do so, we plotted response errors in the current trial as a function of the relative stimulus difference between the previous and current trials (N−1 analysis; Fig. 2a). We quantified the influence of the previous stimulus as the mean sign-adjusted response error using an error-folding procedure (see Methods). To confirm that the observed effect reflected the genuine sequential dependency rather than non-sequential biases in the dataset, we performed an N+1 control analysis (Fig. 2b), in which response errors in the current trial were related to stimuli presented in the subsequent trial. The sign-adjusted response error in the N+1 analysis (–0.01 ± 0.07 mm) was not significantly different from zero (*t*(22) = –0.10, *p* = 0.92, Cohen’s d = –0.02), indicating the absence of systematic non-sequential bias. In contrast, the sign-adjusted response error in the N–1 analysis (0.27 ± 0.08 mm) was significantly larger than that in the N+1 analysis (*t*(22) = 3.06, *p* = 5.7 10^-3^, Cohen’s d = 0.64; Fig. 2c). These results demonstrate that perception of the current slope was systematically attracted toward the immediately preceding one.

**Figure 2.**
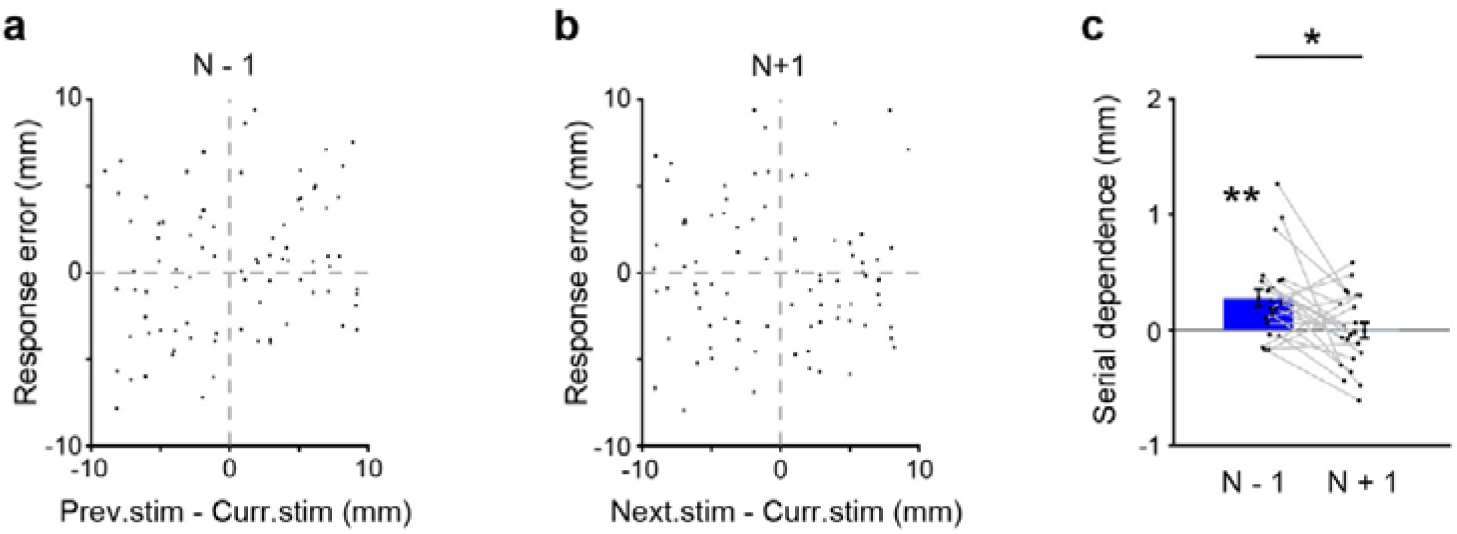
Serial dependence in haptic perception during manual exploration. **a** Response error (reported slope height minus presented slope height) in the current trial plotted as a function of the difference in presented slope height between the previous and current trials for a representative participant. **b** N+1 control analysis. Response errors were plotted as in (**a**). **c** Magnitude of serial dependence quantified as the sign-adjusted response error in the N−1 (blue) and N+1 (light blue) analyses (n = 23 participants). Data are presented as mean ± SEM. \**p* < 0.05, paired two-tailed t-test. \*\**p* < 0.05, two-tailed one-sample t-test.

We next asked how far back in the trial sequence this attractive bias extended. To address this question, we calculated sign-adjusted response errors for stimuli presented one to six trials before the current trial and compared each with the N+1 control. Only the stimulus presented one trial earlier (1-back) produced a significant attractive bias (Fig. 3), whereas stimuli presented further back in the sequence had no detectable effect. Consistent with the time window over which serial dependence emerges (Fischer and Whitney, 2014; Fritsche et al., 2020; Manassi et al., 2023), the present experiment showed serial dependence across trials, with a mean inter-trial interval of 8.24 ± 0.91 s (mean ± SD).

**Figure 3.**
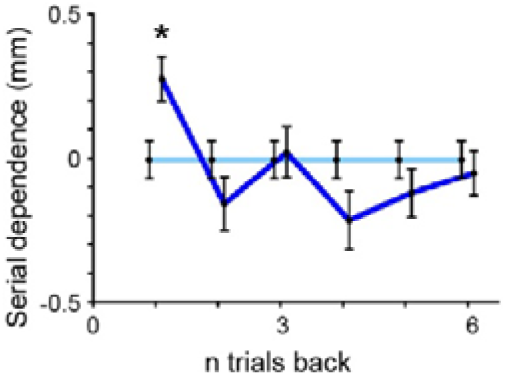
Temporal extent of serial dependence in haptic perception. Magnitude of serial dependence, quantified as the sign-adjusted response error, for stimuli presented one to six trials before the current trial (blue) and for the corresponding N+1 control analyses (light blue) (n = 23 participants). Data are presented as mean ± SEM. \**p* < 0.05 (paired two-tailed *t*-test with Bonferroni correction).

Finally, to verify that the results were not an artifact of the trial-balancing procedure, we repeated all analyses using the complete dataset without trial balancing, while applying the same correction procedures for central tendency, overall bias and edge effects. The results were qualitatively identical (see Supplementary Materials).

### No evidence of motor serial dependence during active touch

The analysis above showed that haptic perception of slope height was systematically biased toward the previous stimulus. However, this effect could have arisen from behavioral changes rather than perceptual processing itself. To examine this possibility, we quantified five kinematic parameters that could serve as a clue for judging slope height: (1) mean handle height while moving along the slope, (2) mean handle height around the target position, (3) total force applied to the handle while moving along the slope, (4) duration of movement along the slope, and (5) movement speed along the slope.

Because no predefined expected value exists for each kinematic parameter at a given stimulus height, we calculated kinematic deviations relative to a stimulus-specific baseline estimated separately for each participant and parameter using linear regression (see Methods). We then examined whether the previous stimulus systematically influenced these deviations. Specifically, we analyzed the relationship between deviations in each kinematic parameter on the current trial and the relative stimulus difference between the previous and current trials (N–1 analysis; Fig. 4). We quantified the influence of the previous stimulus by calculating sign-adjusted kinematic deviation (see Methods).

**Figure 4.**
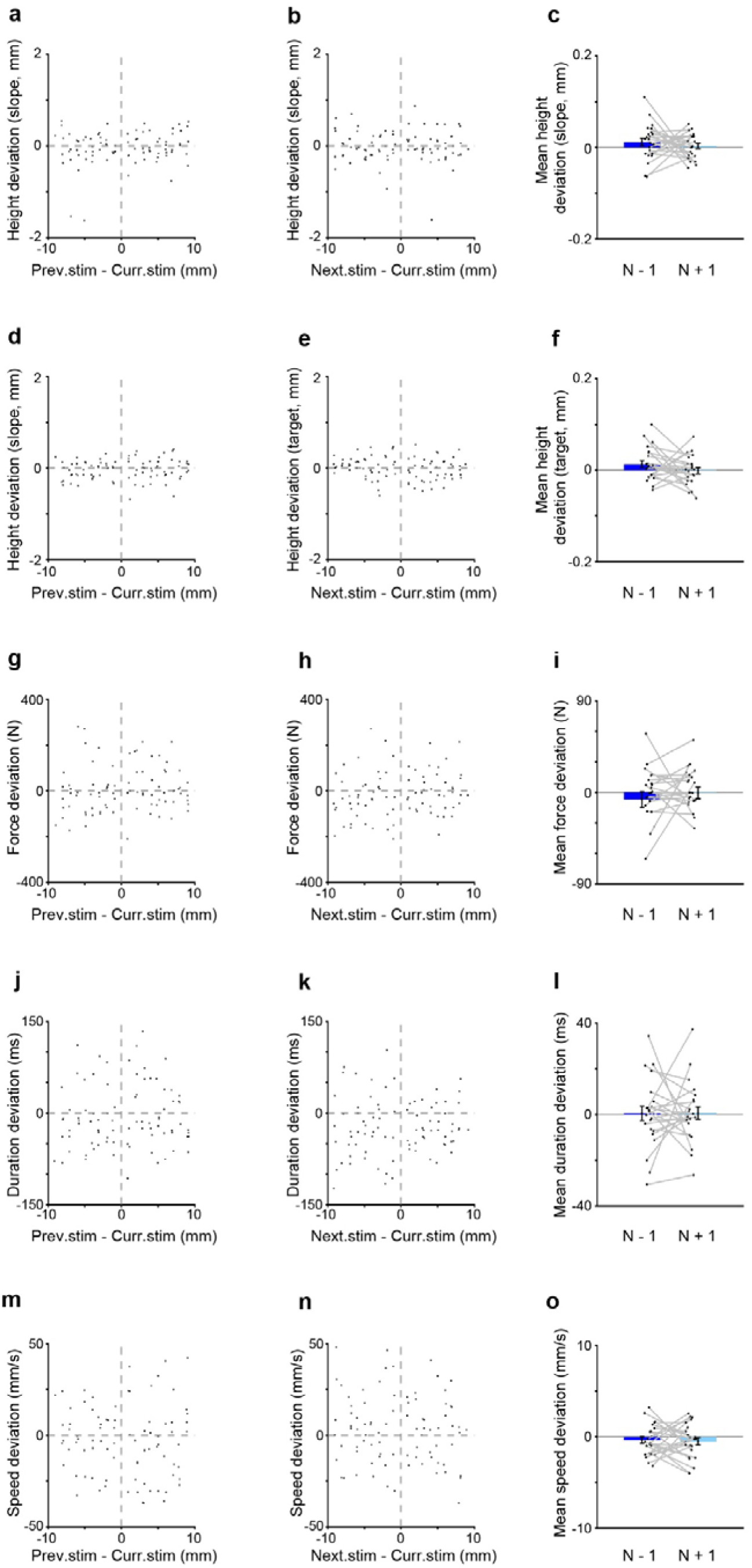
No systematic effects of previous stimuli on exploratory hand movements. **a, d, g, j, m** Kinematic deviations in the current trial plotted as a function of the difference in presented slope height between the previous and current trials (N−1 analysis) for a representative participant. Positive values indicate deviations consistent with attraction toward the previous stimulus. The analyzed kinematic parameters were **(a)** mean handle height while moving along the slope, **(d)** mean handle height around the target position, **(g)** total force applied to the handle while moving along the slope, **(j)** duration of movement along the slope, and **(m)** movement speed along the slope. **b, e, h, k, n** Corresponding N+1 control analyses, in which kinematic deviations in the current trial were plotted as a function of the difference in presented slope height between the current and subsequent trials. **c, f, i, l, o** Mean sign-adjusted kinematic deviations across participants (n = 23) for the N−1 analysis (blue) and the N+1 control analysis (light blue). Sign adjustment was performed so that positive values indicate attraction toward the previous stimulus and negative values indicate repulsion. None of the five kinematic parameters exhibited significant effects of previous stimuli. Data are presented as mean ± SEM. Statistical significance was assessed using two-tailed one-sample *t*-tests and paired two-tailed *t*-tests, as appropriate.

None of the five kinematic parameters showed a significant effect of the previous stimulus (height during movement, *t*(22) = 1.43, *p* = 0.17, Cohen’s d = 0.30, Fig. 4c; height at the target, *t*(22) = 1.81, *p* = 0.08, Cohen’s d = 0.38, Fig. 4f; force, *t*(22) = –0.90, *p* = 0.38, Cohen’s d = –0.19, Fig. 4i; duration, *t*(22) = 0.15, *p* = 0.88, Cohen’s d = 0.03, Fig. 4l; speed, *t*(22) = –1.03, *p* = 0.31, Cohen’s d = –0.21, Fig. 4o). Thus, although perceptual judgments exhibited robust serial dependence, none of the five kinematic parameters showed a systematic effect of the previous stimulus. We obtained similar results when we analyzed the data without a trial-balancing procedure (Supplementary Fig. S1).

We further examined whether perceptual error was associated with deviation in the five kinematic parameters during active touch. For each participant, we calculated the correlation coefficients between perceptual errors and deviations in each of the five kinematic parameters. Correlations were weak across participants for all five parameters (mean ± SD: height during movement, 0.04 ± 0.14; height at the target, 0.08 ± 0.19; force, 0.04 ± 0.15; movement duration, 0.07 ± 0.16; movement speed, 0.09 ± 0.16; Fig. 5). Together, these results indicate that trial-to-trial variations in perceptual errors were not systematically associated with variations in the quantified kinematics.

**Figure 5.**
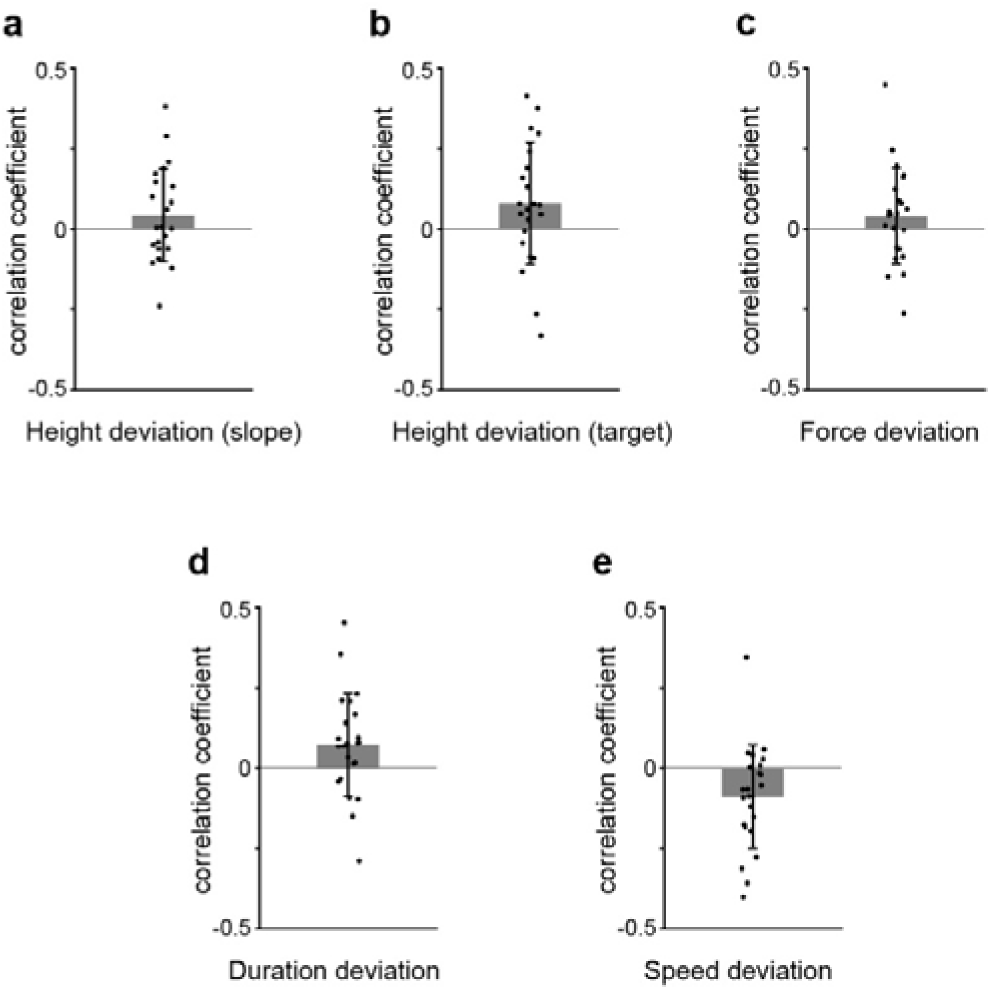
Correlations between perceptual errors and deviations in quantified kinematics. Pearson’s correlation coefficients between perceptual errors and deviations in **(a)** mean handle height while moving along the slope, **(b)** mean handle height around the target position, **(c)** total force applied to the handle while moving along the slope, **(d)** duration of movement along the slope, and **(e)** movement speed along the slope. Correlation coefficients were calculated separately for each participant across trials. Bars and error bars indicate the mean and SD across participants, respectively, and individual dots represent individual participants (n = 23). The horizontal line indicates zero correlation.

## Discussion

In this study, we developed a novel device to quantitatively assess haptic perception during active manual exploration and used it to investigate sequential effects in haptic perception. Participants estimated the height of randomly presented virtual slopes while we simultaneously recorded kinematics of hand movements. We found that haptic perception exhibited serial dependence, consistent with previous observations in the visual, auditory, and olfactory domains. Specifically, perception of the current slope was systematically biased toward the previously explored slope. By contrast, none of the quantified kinematic parameters showed corresponding sequential effects. These findings show that serial dependence extends to active haptic perception and suggest that the quantified kinematics are unlikely to explain the observed perceptual bias.

In this study, we presented virtual slopes with heights ranging from 1 to 20 mm in 1 mm increments in random order. We determined this stimulus range based on the manipulandum’s mechanical limitations, including its range of motion and maximum force output. Using the error-folding approach, we estimated the magnitude of serial dependence to be 0.27 mm, indicating that the expected perceptual bias was substantially smaller than the full stimulus range. Furthermore, participants exhibited a mean discrimination threshold of 3.69 mm (Supplementary Methods and Fig. S2). Because the stimulus spacing (1 mm) was considerably finer than the discrimination threshold, the stimulus set provided sufficient resolution to detect subtle trial-by-trial perceptual biases. Together, these findings indicate that the selected stimulus range and resolution were appropriate for quantifying serial dependence in haptic perception.

The attractive influence of the previous stimulus observed in the present study contrasts with classical tactile aftereffects, which typically induce repulsive perceptual biases. For instance, prolonged exploration of a curved surface causes a subsequently explored flat surface to be perceived as curved in the opposite direction, a phenomenon known as the curvature aftereffect (van der Horst et al., 2008; Vogels et al., 1997, 1996). Unlike serial dependence, the haptic aftereffect generally requires sustained stimulus exposure and typically emerges after at least 2 s of continuous contact. In the present experiment, however, participants contacted the inclined position of the virtual slope for only approximately 0.5 s. Such brief exposure is unlikely to induce substantial adaptation-related aftereffects. The different stimulus durations required to observe repulsive adaptation and attractive serial dependence across experiments may reflect complementary computational strategies rather than conflicting mechanisms. As time elapses, the environment becomes increasingly likely to change, inducing a tendency for the perceptual system to enhance sensitivity to novel inputs by reducing responses to recently experienced input. By contrast, over shorter timescales after a stimulus, the environment is more likely to remain stable, so the brain tends to improve perceptual stability by incorporating recent sensory experience into current input. The experimental conditions used in this study may have minimized adaptation, allowing serial dependence to emerge as the dominant history-dependent influence on perceptual judgments (Moon and Kwon, 2022; Sheehan and Serences, 2022).

In contrast to our findings, previous studies showed that serial dependence can influence eye movements during sensory sampling. Goettker and Stewart demonstrated that oculomotor serial dependence is driven primarily by retinal error signals (Goettker and Stewart, 2022). One possible explanation for this difference lies in how tightly sensory information constrains motor behavior. In smooth pursuit, retinal motion is mapped relatively directly onto a comparatively constrained oculomotor response required to maintain target tracking. In active haptic exploration, by contrast, object properties can be acquired through multiple combinations of movement trajectory, force, speed, and duration. A fundamental requirement of haptic exploration is that object features can be extracted across different sequences of exploratory movements, while the movements themselves are flexibly selected according to sensory information, task demands, and behavioral goals (Hartmann, 2009).

Furthermore, exploratory movements generate mechanical interactions with the environment that themselves constitute sensory input (Zweifel and Hartmann, 2020). Ongoing tactile and proprioceptive feedback from the current object can therefore continuously shape exploratory movements as sensory information is acquired. Previous haptic information may thus influence exploration within a flexible motor process that is continuously updated by current sensory input, although serial dependence was not evident in the movement kinematics.

Serial dependence has been reported across multiple sensory modalities, including vision, audition, olfaction, and haptic perception, despite substantial differences in how sensory information is acquired and represented across these modalities (Burg et al., 2022; Fischer and Whitney, 2014; Motala et al., 2020). This cross-modal phenomenon is compatible with hierarchical accounts of serial dependence, in which perceptual history is integrated at relatively high levels to form priors that are subsequently fed back to modulate modality-specific sensory representations at lower levels, providing a potential common framework for history-dependent biases across modalities (Cicchini et al., 2024). Working memory may provide one substrate for maintaining such higher-level representations, allowing recently acquired information to persist beyond the immediate sensory input and remain available for subsequent perceptual processing (Baddeley, 2012; Markov et al., 2024). The present finding of haptic serial dependence raises the possibility that recently acquired haptic information may likewise be retained at higher representational levels and integrated with sensory information acquired from the current object.

A few limitations should be addressed. According to previous studies, perceptual judgments are influenced by history-dependent processes operating over multiple trials, reflecting continuously updated beliefs about the statistical structure of the environment (de Lange et al., 2018; Petzschner et al., 2015). Although we removed a central-tendency bias, it cannot completely dissociate slowly varying Bayesian-like influences from previous sensory experiences. We therefore do not claim that the observed bias represents only “serial” dependence. Nevertheless, our results indicate that short-term serial dependence, typically from the previous perception to the current one, was the dominant perception history-dependent effect. Specifically, the sign-adjusted response error was significantly smaller in the N−2 analysis than in the N−1 analysis (Fig. 3b), consistent with the characteristic temporal profile of serial dependence (Manassi et al., 2023). Because any bias observed in the N−2 analysis necessarily includes serial dependence propagated through the intervening N−1 trial, the contribution of slower Bayesian-like history effects is likely to be relatively small. Together, these findings suggest that, after accounting for central tendency, perceptual judgments were primarily shaped by short-term serial dependence. Haptic perception of object geometry is constructed by integrating multiple sources of sensory information acquired during active touch. In particular, cutaneous afferents provide information about local surface properties, whereas proprioceptive signals from muscles, joints, and tendons convey information about hand position and movement (Berryman et al., 2006; Hsiao, 2008; Sobinov and Bensmaia, 2021; Voisin et al., 2002). These sensory signals are integrated with motor commands to construct a representation of object geometry, including both local tactile features and global properties such as size and shape (Delhaye et al., 2018). Although the present findings suggest that serial dependence arises primarily from perceptual processing rather than from the quantified kinematics, the neural mechanisms underlying this perceptual bias remain unclear. Disentangling the respective contributions of cutaneous and proprioceptive signals during active touch remains experimentally challenging in humans and represents an important direction for future research.

In conclusion, we demonstrated attractive serial dependence in haptic perception of object geometry. Systematic changes in hand movements associated with active touch did not explain haptic serial dependence. Our findings extend the concept of serial dependence to haptic perception in which sensory information is acquired through hand movements.

## Author contributions

Conceptualization: TU. Investigation: YZ. Data Curation: YZ. Formal Analysis: YZ, TU. Methodology: YZ, TN, TU. Resources: TU. Software: YZ, TN, TU. Project Administration: TU. Supervision: TH, TU. Validation: TU. Visualization: YZ, TU. Writing – Original Draft Preparation: YZ, TU. Writing – Review & Editing: YZ, TN, TH, TU. Funding Acquisition: TU.

## Data availability

The data supporting the findings of this study are available in the Zenodo repository at https://doi.org/10.5281/zenodo.13958815.

## Code availability

Analysis scripts are available in the Zenodo repository at https://doi.org/10.5281/zenodo.13958815.

## Declaration of generative AI and AI-assisted technologies in the manuscript preparation process

During the preparation of this work, the authors used ChatGPT (OpenAI) in order to improve the language, readability, and organization of the manuscript. After using this tool, the authors reviewed and edited the content as needed and take full responsibility for the content of the publication.

