## Supplementary Material for "Serial dependence in haptic perception during active manual exploration"

Supplementary Materials for

**Serial dependence in haptic perception**

**during active manual exploration**

Yunshan Zhang *et al.*

**This PDF file includes:**

Supplementary Results

Figs. S1-2

**Supplementary Results**

Robustness analysis without trial subsampling

To evaluate whether the trial subsampling procedure influenced the results, we repeated all analyses using the same stimulus range (5–16 mm) but without trial subsampling. Consistent with the main analyses, the sign-adjusted response error in the N–1 analysis (0.27 ± 0.05 mm) was significantly greater than zero (*t*(22) = 5.52, *p* = 1.52 ×10^-5^, Cohen’s *d* = 1.15; Fig. S1a) and significantly larger than that in the N+1 analysis (-0.01 ± 0.05 mm; *t*(22) = 4.31, *p* = 2.86 ×10^-4^, Cohen’s *d* = 0.90; Fig. S1a). The N+1 error did not differ from zero (*t*(22) = -0.17, *p* = 0.86, Cohen’s *d* = -0.04).

For the movement analysis, only the mean handle height around the target position (0.01 ± 0.00 mm) showed a positive deviation from zero in the N−1 analysis (*t*(22) = 2.87, *p* = 0.01, Cohen’s d = 0.59). However, this effect did not differ significantly from the corresponding N+1 control analysis (*t*(22) = 1.70, *p* = 0.10, Cohen’s d = 0.36), indicating a lack of temporal specificity. None of the five kinematic parameters therefore showed robust history-dependent biases in exploratory hand movements. These findings indicate that the dissociation between perceptual serial dependence and exploratory movements was not driven by the trial subsampling procedure (Fig. S1).

Measurement of haptic discrimination threshold

After Experiment 1, we conducted a two-alternative forced choice (2AFC) task to measure a discrimination threshold for perceiving the height of the virtual slope in 21 participants (Experiment 2). The experimental setup was similar to that of Experiment 1. However, the participants were requested to move the handle twice consecutively, encountering two slopes of different heights within each trial. The interval between the end of the first sub-trial and the beginning of the second sub-trial was 4.7 s. After completing the two sub-trials, participants were required to indicate whether the slope in the second sub-trial was higher or lower than that in the first sub-trial by pressing a key labeled 'height' or 'low' with their left hand. Experiment 2 consisted of three runs, with 100 trials per run. A run lasted 20 to 30 minutes, with an 8-minute break between runs (Fig. S2a).

To quantify the discrimination threshold in the 2AFC task, the difference in slope height between the first and second slopes was plotted on the x-axis, with y = 1 when the second slope was perceived as higher and y = 0 when it was perceived as lower. The data were then fitted with a logistic regression model. The slope-height differences corresponding to response probabilities of 0.25 and 0.75 were determined from the fitted logistic function, and half of the difference between these values was defined as the just noticeable difference (JND). This analysis yielded a JND of 3.69 ± 1.58 mm (mean ± SD, *n* = 21; Fig. S2b).

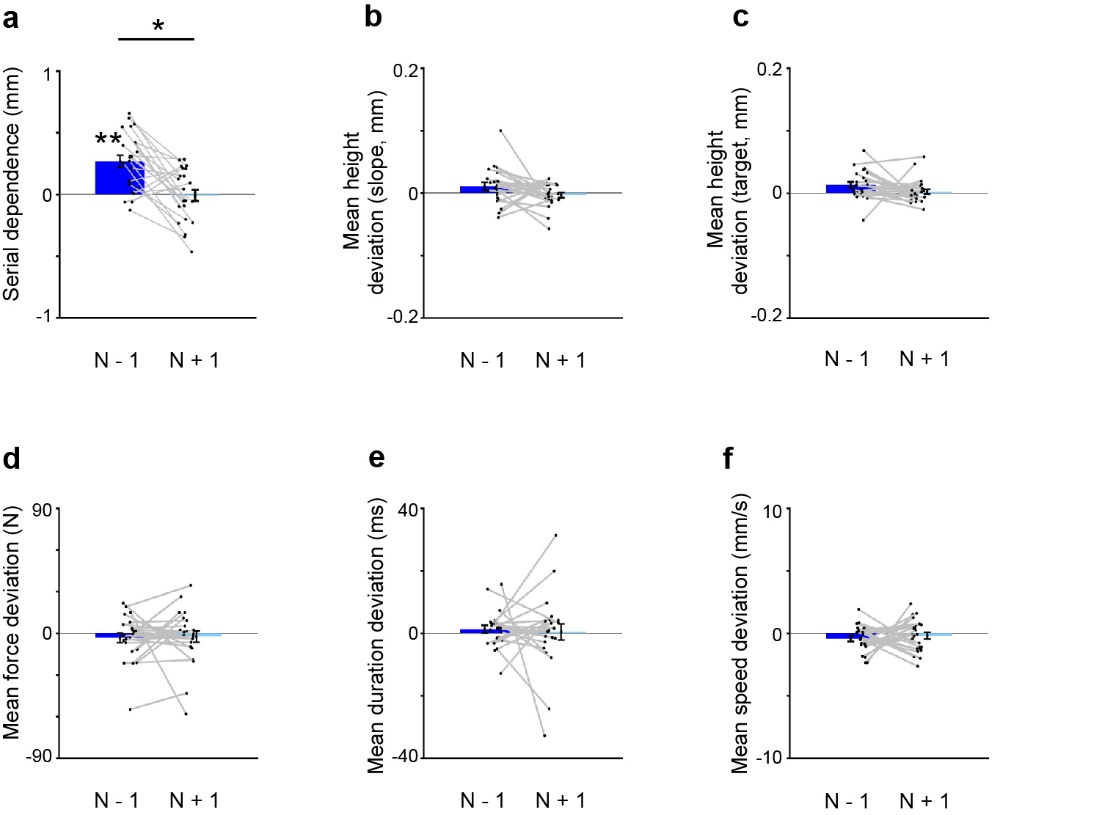

**Figure S1. Serial dependence in height perception and movement kinematics without trial subsampling**

**a** Magnitude of serial dependence (sign-adjusted response error) in the N–1 analysis (blue) and N+1 analysis (light blue) (*n* = 23 participants). **b-f** Mean sign-adjusted kinematic deviations across participants (n = 23) for the N−1 analysis (blue) and N+1 control analysis (light blue). In a-f, results are calculated from data without trial subsampling. Data are presented as mean ± SEM. *P < 0.05, paired two-tailed t-test. **P < 0.05, two-tailed one-sample t-test.

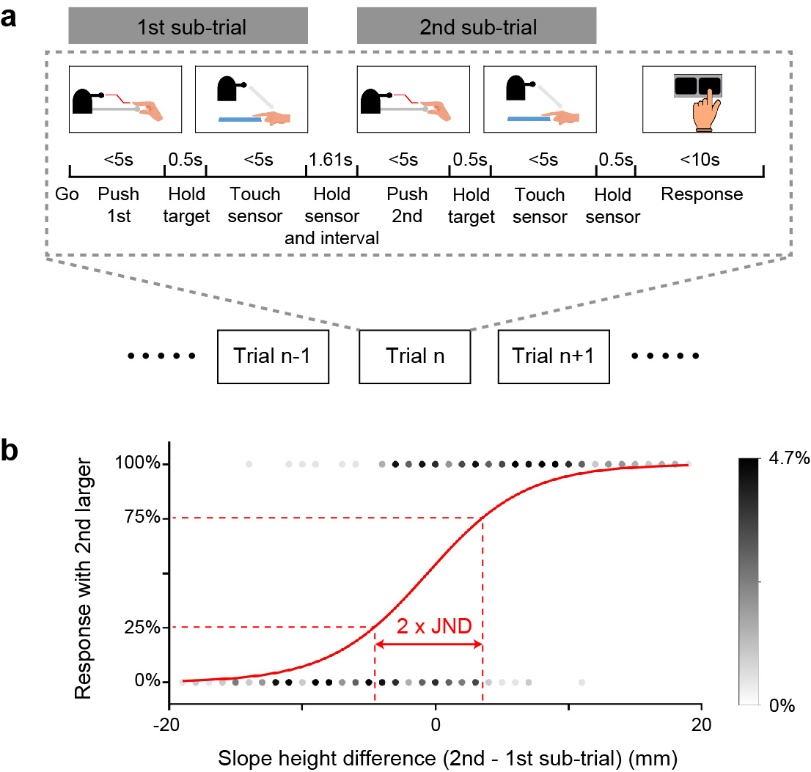

**Figure S2. Two-alternative forced-choice (2AFC) experiment and estimation of the just noticeable difference (JND).**

**a** Experimental event sequence in 2AFC experiment. Each trial consisted of two consecutive sub-trials. Response was not required in the first sub-trial. Participants compared the heights of the slopes between the first and second sub-trials by pressing one of two buttons representing high or low. **b** Relationship between the difference in slope height between the first and second sub-trials and the proportion of responses indicating that the second slope was perceived as higher for one participant. The grayscale indicates the proportion of trials represented by each data point. A red line represents the fitted logistic regression curve. Half of the difference between the slope-height values corresponding to response probabilities of 25% and 75% on the fitted logistic curve was defined as the JND.
